# Early zygotic gene product Dunk interacts with anillin to regulate Myosin II during *Drosophila* cleavage

**DOI:** 10.1101/2022.02.14.480462

**Authors:** Jiayang Chen, Bing He

## Abstract

*Drosophila* cellularization is a special form of cleavage that converts syncytial embryos into cellular blastoderms by partitioning the peripherally localized nuclei into individual cells. An early event in cellularization is the recruitment of non-muscle myosin II (“myosin”) to the basal tip of cleavage furrows, where myosin forms an interconnected basal array before reorganizing into individual cytokinetic rings. The initial recruitment and organization of basal myosin are regulated by a cellularization-specific gene, *dunk*, but the underlying mechanism is unclear. Through a genome-wide yeast two-hybrid screen, we identified anillin (Scraps in *Drosophila*), a conserved scaffolding protein in cytokinesis, as the primary binding partner of Dunk. We show that Dunk regulates the localization of anillin at the cleavage furrows during early cellularization and functionally interacts with anillin in regulating basal myosin. Furthermore, we show that anillin colocalizes with myosin since the onset of cellularization and is required for the initial recruitment and maintenance of myosin at the basal array, prior to the well-documented function of anillin in regulating cytokinetic ring assembly. Based on these results, we propose that Dunk regulates myosin recruitment and organization during early cellularization by interacting with and regulating anillin.

## Introduction

Cytokinesis, the process that partitions the cellular contents of one dividing cell into two daughter cells, is one of the most critical events for live organisms (Green et al., 2012; Rappaport, 1971). As cytokinesis starts, a contractile ring composed of F-actin and non-muscle myosin II (hereafter “myosin”) is formed at the cell equator. The subsequent constriction of this ring drives cleavage furrow invagination and eventually pinches off the two daughter cells (Eggert et al., 2006; Green et al., 2012). The tight regulation of myosin activity, organization, and localization enables the proper constriction and the separation of the daughter cells (Green et al., 2012; Pollard and O’Shaughnessy, 2019). In this study, we sought to understand how cells make use of the conserved cytokinesis machinery to achieve a special form of cleavage.

*Drosophila* cellularization is atypical cytokinesis that mediates cellular blastoderm formation during early *Drosophila* embryogenesis (Mazumdar and Mazumdar, 2002). Before cellularization, the *Drosophila* embryo undergoes 13 rounds of nuclear division without cytokinesis, generating a syncytial blastoderm with ∼ 6,000 nuclei aligned near the surface of the embryo (Foe and Alberts, 1983). At the onset of cellularization, the plasma membrane begins to invaginate and form cleavage furrows between the syncytial nuclei. Approximately an hour later, a monolayer of epithelial cells is formed at the peripheral region of the embryo, marking the end of cellularization (Mazumdar and Mazumdar, 2002). Similar to conventional cytokinesis, active myosin is recruited to the cell cortex at the onset of the cellularization and becomes concentrated at the leading edge of nascent cleavage furrows, also known as “cellularization front” or “furrow canals” (Fig. 1A) (Fullilove and Jacobson, 1971; Royou et al., 2004). This initial recruitment of myosin is associated with a cortical flow of punctum-like apical myosin structures towards the nascent furrows (He et al., 2016). This initial, flow-associated recruitment is followed by direct recruitment of myosin from the cytoplasm to the furrow tip (Royou et al., 2004; He et al., 2016). Myosin recruited to the cellularization front first organizes into an interconnected hexagonal array that resembles the spatial pattern of the ingressing furrows. Approximately 30 minutes into cellularization, this basal myosin array reorganizes into individual myosin rings, which contract and eventually close off the base of the newly formed cells (Fig. 1A) (Royou et al., 2004; Xue and Sokac, 2016). In addition to myosin, many other structural components and regulators of the cytokinetic ring are shared between cellularization and conventional animal cytokinesis, such as actin (Schejter and Wieschaus, 1993), RhoA (Rho1 in *Drosophila*) and its regulators (Crawford et al., 1998; Padash Barmchi et al., 2005; Grosshans et al., 2005; Wenzl et al., 2010; Mason et al., 2016; Sharma and Rikhy, 2021), anillin (Scraps in *Drosophila*) (Field et al., 2005; Field and Alberts, 1995), septin (Adam et al., 2000; Field et al., 2005), and the formin protein Diaphanous (Afshar et al., 2000; Grosshans et al., 2005). These conserved cytokinetic components are mainly maternally provided, many of which are involved in regulating the localization and function of myosin during cellularization.

**Figure 1.**
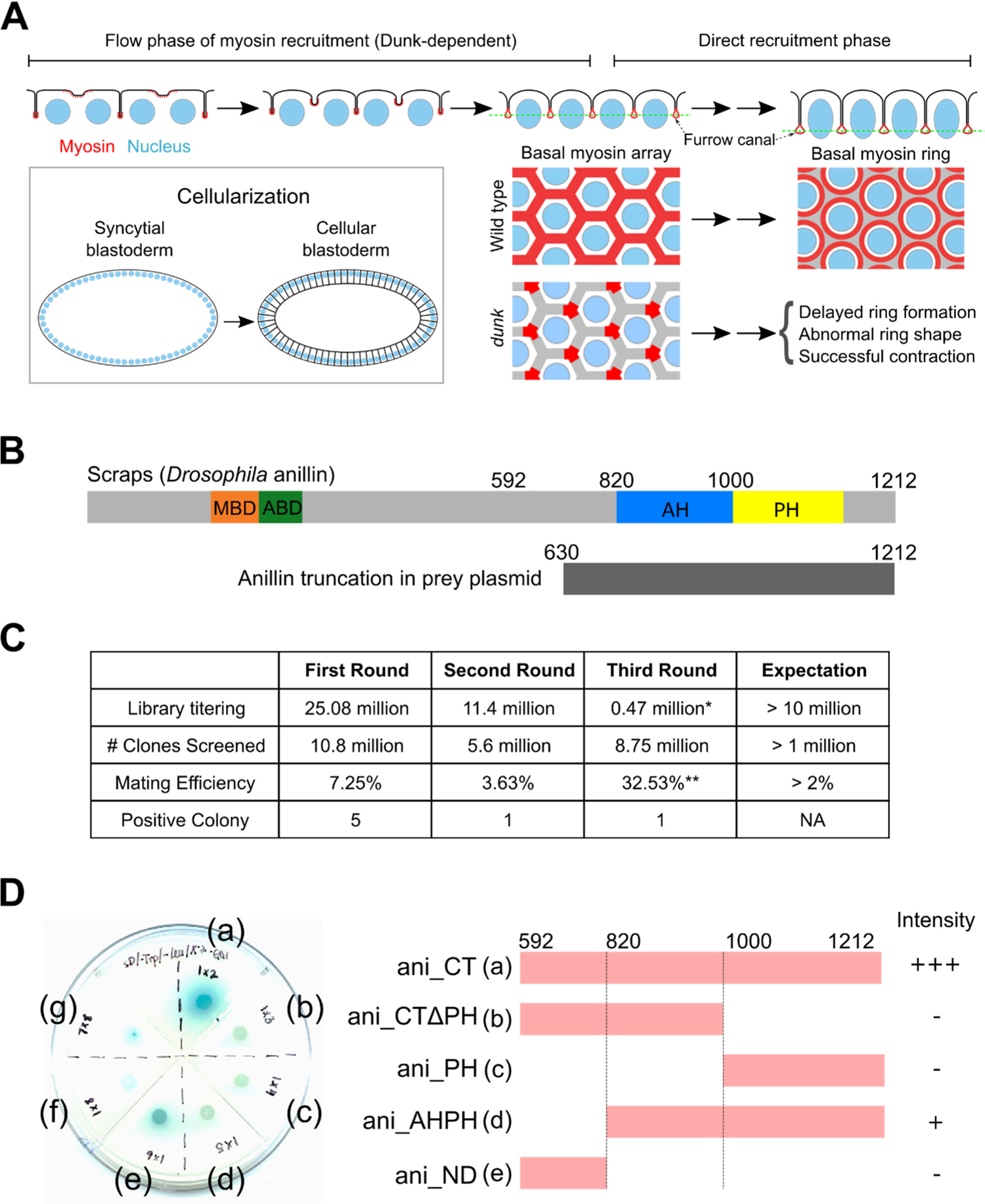
Dunk interacts with anillin through its C-terminus. **(A)** Schematics showing myosin recruitment to the cleavage furrows during *Drosophila* cellularization and the basal myosin phenotype in *dunk* mutant embryos. **(B)** Top: domain structure of Scraps, the *Drosophila* anillin. The C-terminal region of anillin is the sole Dunk-interactor identified in our yeast-two-hybrid screen. Bottom: The schema of the truncated anillin protein encoded by the anillin sequence in the prey library. The information of anillin truncation was obtained by sequencing the prey plasmids from the positive colonies. All seven positive colonies were sequenced, and all started around 630 a.a. of the full length anillin. **(C)** A table showing the library titering, numbers of clones screened, mating efficiency, and the number of the positive clones for each repeat. The library titering indicates whether the library is healthy enough and contains enough yeast cells; the number of clones screened is the indicator of the number of yeast cells from the library that have been gone through in each screen; mating efficiency indicates whether bait cell and prey cell mated successfully (Mate & Plate™ Library - Universal *Drosophila* (Normalized), Takara, 630485). The unexpected and abnormal titering (*) and matting efficiency (**) of the third round of repeats were resulted from the low number of colonies growing on the SD/-Leu control plate (See Materials and Methods for details). **(D)** A schematic outlining the constructs used for pair-wise yeast two-hybrid assay. One representative raw SD/–Leu/–Trp/X-alpha-Gal/AbA agar plate showing the colonies for each pair of bait and prey. (a): full length C-terminal anillin (ani_CT), 592-1212 a.a.; (b): AH domain (ani_CTΔPH), 592-1000 a.a.; (c): PH domain (ani_PH), 1000-1212 a.a.; (d): AH domain and PH domain (ani_AHPH), 820-1212 a.a.; (e): C-terminal region without AH domain and PH domain (ani_ND), 592-820 a.a.; (f): negative control; (g): positive control. Only the prey constructs containing the entire C-terminal region (“ani_CT”) or both the AH and PH domains (“ani_AHPH”) showed positive interaction with Dunk.

In addition to the conserved cytokinetic factors, a small number of “cellularization-specific” genes are also involved in regulating cellularization. Five such “cellularization genes” have been identified: *nullo*, *serendipity-α*, *bottleneck*, *slam* and *dunk* (Merrill et al., 1988; Wieschaus and Sweeton, 1988; Schweisguth et al., 1990; Rose and Wieschaus, 1992; Schejter and Wieschaus, 1993; Lecuit et al., 2002; He et al., 2016). The transition from the syncytial stage to cellularization is coincident with the genome-wide activation of zygotic gene expression in the embryo (Renzis et al., 2007; Liang et al., 2008; Tadros and Lipshitz, 2009). The five cellularization genes are all expressed in a short time window around the onset of cellularization, and the corresponding proteins rapidly disappear during late cellularization. The products of these genes are all localized to the furrow canals and are involved in regulating the basal actin-myosin (actomyosin) structures, either directly or indirectly. Dunk is critical for the flow phase of myosin recruitment during the first ten minutes of cellularization (He et al., 2016). Slam, on the other hand, is important for the subsequent, direct recruitment of myosin to the cellularization front (Lecuit et al., 2002; Wenzl et al., 2010; Acharya et al., 2014; He et al., 2016). Nullo and Serendipity-α promote actin assembly at the furrow canals, supporting actomyosin contractility and preventing degeneration of cleavage furrows (Schweisguth et al., 1990; Rose and Wieschaus, 1992; Sokac and Wieschaus, 2008; Zheng et al., 2013). Finally, Bottleneck acts as an actin-crosslinking protein that functions to dampen actomyosin contractility and restrain the transition from basal myosin network into individual myosin rings, thereby ensuring the proper timing of basal closure (Schejter and Wieschaus, 1993; Reversi et al., 2014; Krueger et al., 2019). While the molecular functions of several cellularization proteins have started to be elucidated, how Dunk regulates myosin during cellularization remains elusive.

Anillin is a highly conserved protein that plays an important role during cytokinesis (Field and Alberts, 1995; Piekny and Maddox, 2010). The structure of anillin is well defined, revealing multiple protein-protein interaction domains conserved throughout species from *Drosophila* to humans (Piekny and Maddox, 2010). Anillin binds to the actomyosin network through its myosin- and actin-binding domains at its N-terminus and links contractile ring to the cell membrane through anillin homology (AH) domain and a Pleckstrin homology (PH) domain at its C-terminus (Fig.1B) (Piekny and Glotzer, 2008; Sun et al., 2015; Budnar et al., 2019). Anillin’s regulators RhoA and PI(4,5)P_2_ binds to the AH domain of anillin and facilitate its cortical localization (Piekny and Glotzer, 2008; Sun et al., 2015; Budnar et al., 2019). The PH domain of anillin does not display affinity to phospholipids but is required for the cortical localization of anillin (Sun et al., 2015). The PH domain of anillin binds to the septin subunit Peanut (Field et al., 2005; Sun et al., 2015). Although the septin cytoskeleton provides important structural functions required for the completion of cytokinesis, it is dispensable for the membrane localization of anillin (Field et al., 2005; Sun et al., 2015). During conventional animal cytokinesis, anillin functions to promote the temporal and spatial stability of the actomyosin network at the cell equator and facilitate the organization of actomyosin into contractile rings (Piekny and Maddox, 2010). For example, in HeLa cells, loss of anillin leads to destabilization and oscillation of the actomyosin ring around the equatorial region (Piekny and Glotzer, 2008). During *Drosophila* cellularization, loss of anillin leads to disorganization of basal actomyosin rings and failure in basal closure (Thomas and Wieschaus, 2004; Field et al., 2005). In both cases, however, myosin is still recruited to the cleavage furrows and is able to contract (Thomas and Wieschaus, 2004; Field et al., 2005; Piekny and Glotzer, 2008). The lack of obvious phenotype in initial myosin recruitment upon disruption of anillin may be due to functional redundancy, as suggested by studies in cultured cells (Piekny and Glotzer, 2008). In addition to a critical function in cytokinesis, recent studies have also revealed anillin’s function in regulating myosin localization at the cell-cell junctions (Wang et al., 2015) and organizing medial-apical actomyosin network in non-dividing epithelial cells (Arnold et al., 2019). Interestingly, a recent study suggests that anillin may function as an unconventional RhoA effector that feeds back and boosts RhoA’s activity by kinetically increasing its residence time at PI(4,5)P_2_-enriched membrane (Budnar et al., 2019). How the functions of anillin in regulating cellular contractility are involved in an early stage of cytokinesis prior to the formation of the cytokinetic ring is not fully understood.

Here, we report the identification of anillin as a Dunk-binding partner in a genome-wide yeast two-hybrid screen. We provide evidence that Dunk and anillin function together to regulate cortical myosin in early cellularization. Anillin colocalizes with myosin at the cellularization front throughout cellularization, both before and after the basal myosin array reorganizes into contractile rings. In *anillin* mutant, basal myosin is frequently depleted from the edges within the basal array, causing an enrichment of myosin at the vertices that closely resembles the basal myosin phenotype in *dunk* mutant. In the absence of Dunk, anillin and the septin subunit Peanut show aberrant localization at the basal array. Furthermore, *dunk* and *anillin* genetically interact with each other and show similar synthetic effects when combined with loss of Bottleneck. Taken together, we propose that Dunk regulates basal myosin during early cellularization by interacting with anillin and regulating its localization and/or activity.

## Result

### Identification of anillin as a binding partner for Dunk

Previous work has shown that the zygotic gene *dunk* functions to facilitate the flow phase of myosin recruitment during early cellularization. In *dunk* mutant embryos, myosin is progressively depleted from the cellularization front during the first 5 – 10 minutes of cellularization, resulting in a fragmented basal myosin array with a low level of myosin at most edges of the hexagonal array (Fig. 1A) (He et al., 2016). Dunk is a small protein (246 a.a.) with no previously identified homolog or well-characterized structure motifs. In order to understand how Dunk regulates myosin in cellularization, we performed a genome-wide yeast two-hybrid screen to identify Dunk binding partners. We used full-length Dunk protein as the bait to screen a normalized *Drosophila* cDNA library (Mate & Plate™ Library - Universal *Drosophila* (Normalized), Takara, 630485). This library was prepared from a mix of equal quantities of poly-A+ RNAs isolated from the embryo, larva, and adult stage *Drosophila* and covers most of the expressed genes. Scraps, the *Drosophila* homolog of anillin, was the only protein that displayed positive interaction with Dunk in the three repeats of the screen (Fig. 1B, C). Of note, the cDNAs in the library on average cover ≤ 600 amino acids from the C-terminus of the encoded prey proteins (Pretransformed Mate & Plate™ Libraries (January 2008) Clontechniques XXIV(1):26– 27). For example, it only contains the last 583 amino acids (630 - 1212 a.a.) from the C-terminus of anillin (Fig. 1B). It is, therefore, possible that we have missed the binding partner of Dunk that is longer than 600 amino acids and requires its N-terminal portion to bind to Dunk. On the other hand, the result that anillin is the only binding protein identified in the screen suggests that the bait protein in the assay was not “sticky”. Therefore, the interaction we detected was unlikely due to non-specific binding.

Anillin is a multi-domain scaffold protein that binds to the actomyosin network through its N-terminal myosin and actin-binding domains and to the cell membrane through its C-terminal domains (Piekny and Maddox, 2010) (Fig. 1B). The C-terminal portion of Anillin is important for its cortical localization and is well conserved from *Drosophila* to humans (Field et al., 2005; Piekny and Maddox, 2010; Sun et al., 2015). The positive clones identified in our screen all contain the C-terminal portion of anillin, which includes the conserved AH and PH domains (Fig. 1A). However, our attempts to further confirm this interaction by co-immunoprecipitation or *in vitro* protein binding assays were unsuccessful due to various technical challenges. Thus, it remains to be addressed whether Dunk and anillin physically interact with each other.

Given the technical challenges of testing the Dunk-anillin interaction using alternative approaches, we sought to use yeast two-hybrid to confirm the result from the screen and further determine the minimal region of anillin that binds to Dunk. To this end, we generated prey constructs of different truncations of anillin and tested for the interaction with Dunk as the bait in a pairwise yeast two-hybrid assay (Fig. 1D). Only the prey construct containing the entire C-terminal region (“ani_CT”, 592-1212 amino acids) and a truncated fragment containing both the AH and PH domains (“ani_AHPH”, 820-1212 amino acids) showed positive interaction with Dunk. Neither the AH domain nor PH domain alone was sufficient to mediate the interaction (Fig. 1D). Interestingly, the ani_CT construct showed a stronger signal compared to the ani_AHPH construct (Fig. 1D), suggesting that the sequence outside of the AH and PH domains also contributes to this binding. Together, the results of pairwise yeast two-hybrid assay confirmed the finding from the genome-wide screen and further mapped the Dunk binding site on anillin to a region encompassing the conserved AH and PH domains. Because anillin interacts with actomyosin and is a well-known regulator for cytokinesis, we hypothesized that Dunk regulates the formation of basal myosin array through anillin.

### Dunk regulates the spatial distribution of Anillin at the furrow canals

To test our hypothesis, we first asked whether Dunk regulates the localization of anillin during cellularization. Previous studies have shown that anillin is concentrated at the cellularization front and colocalizes with the basal actomyosin rings (Field et. al, 2005). Analysis of anillin localization in wildtype and *dunk* mutant embryos by immunostaining revealed that in *dunk* mutant embryos, anillin was still recruited to the furrow canals, but the spatial distribution of anillin at the furrow canals was abnormal (Fig. 2A). During early cellularization, anillin was evenly distributed across the cellularization front in wildtype embryos, forming an interconnected hexagonal array that was analogous to the basal myosin array. In *dunk* mutant embryos, however, the distribution of anillin at the basal array was less homogeneous compared to the wildtype (Fig. 2A, red and green arrows showing enrichment and depletion of the anillin signal, respectively). By the time anillin started to reorganize into individual rings in wildtype embryos (Fig. 2A, Early-mid cellularization), anillin signal in *dunk* mutant embryos became more homogeneous across the leading edge, suggesting a partial recovery from the early defect (Fig. 2A, Early-mid cellularization). However, in contrast to the wildtype embryo where anillin was enriched in curved bundles that resemble part of the cytokinetic rings (Fig. 2A, yellow arrows), there was no sign of contractile ring formation in *dunk* mutant embryos (Fig. 2A, cyan arrows). By the time anillin clearly formed discrete rings in wildtype embryos (Fig. 2A, “Mid cellularization” and “Mid-late cellularization”, yellow arrowheads), anillin in *dunk* mutant appeared to just start to rearrange into individual rings. The shape of these rings was irregular, and the neighboring rings were often not well resolved (Fig. 2A, cyan arrowheads). Finally, after the basal rings were formed in *dunk* mutant embryos, they were able to contract and became smaller over time (Fig. 2A, “Late cellularization”). The apparent delay in the ring formation, the abnormal ring morphology, as well as the ability for the ring to contract are all analogous to the basal myosin phenotype in *dunk* mutant embryos (He et al., 2016). The septin subunit Peanut, which depends on anillin to be recruited to the furrow canals during cellularization (Field et. al, 2005), showed similar localization defects as anillin in *dunk* mutant embryos (Fig. 2B). Together, these results indicate that while Dunk is dispensable for the recruitment of anillin and septin to the cellularization front, it regulates the spatial organization of anillin and septin across the basal array and facilitates the rearrangement of these proteins into ring configurations.

**Figure 2.**
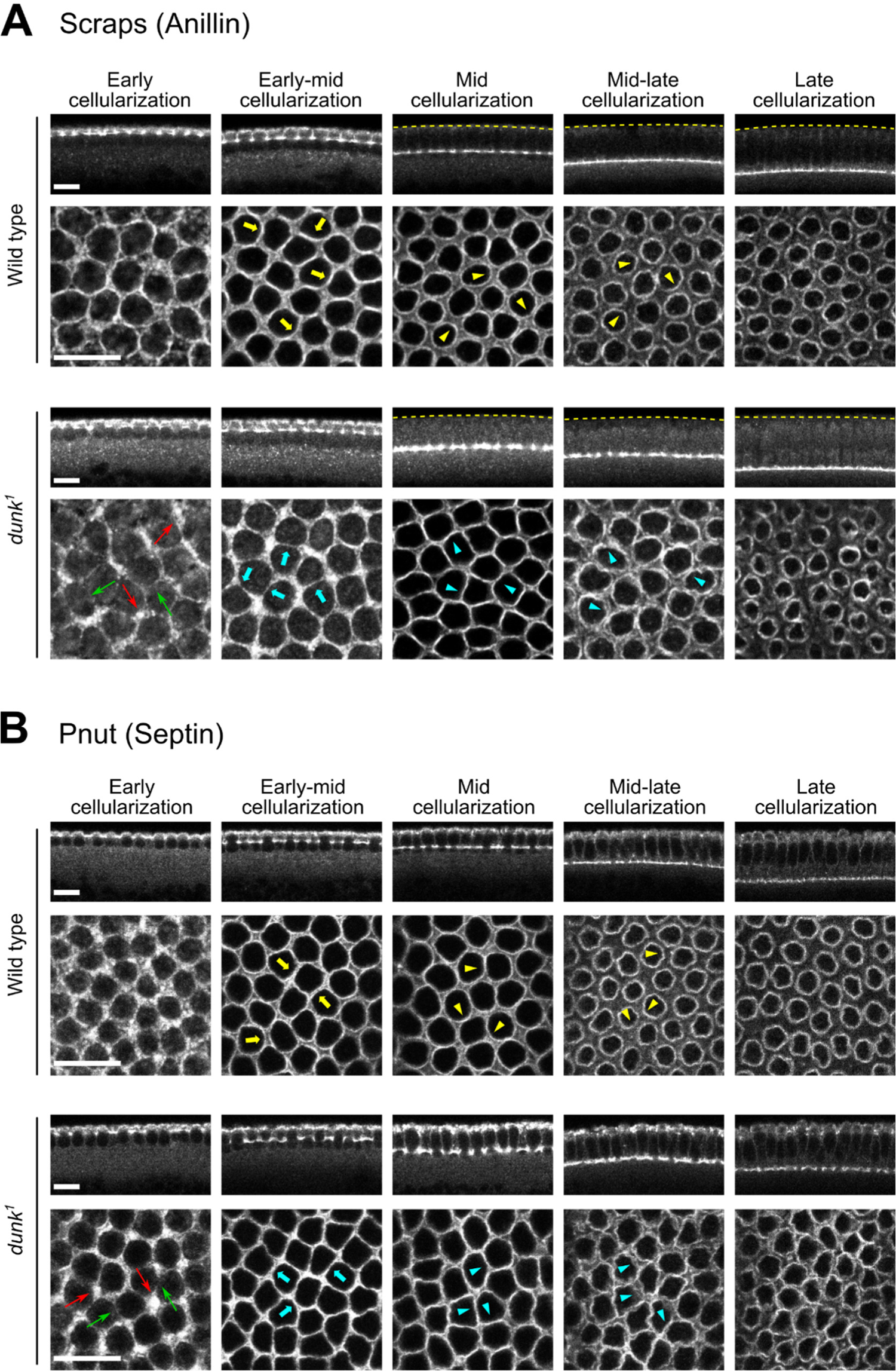
Dunk regulates the localization of anillin and the septin Peanut during cellularization. Anillin (A) and the septin Peanut (Pnut, B) are mislocalized in *dunk^1^* mutant embryos. Immunostaining showing localization of Anillin (Scraps) (A) and Septin (Peanut) (B) in cross-sections (top) and en face sections at the furrow canals (bottom). Anillin and Septin show similar defects in *dunk^1^* mutant embryos. During early cellularization, anillin and Septin are recruited to the furrow canals, but their spatial distribution at the furrow canals appears to be less homogeneous compared to the wildtype controls. The signal is often enriched at the vertices (red arrows) and depleted from the edges (green arrows). During early to mid-cellularization, anillin and septin in the control embryos become enriched at the nascent contractile rings (cyan arrows). In contrast, there is no clear sign of ring formation for anillin or septin in *dunk^1^* mutant embryos at the similar stage. Although anillin and septin in the mutant embryos eventually form individual rings during mid- to late cellularization, the rings are less circular and more irregular (cyan arrowheads) compared to those in the wild type embryos (yellow arrowheads). Yellow dotted lines in A mark the apical surface of the embryo. All scale bars:10 µm.

### Anillin colocalizes with myosin and shows similar cortical dynamics as myosin in early cellularization

The early defects in basal myosin organization in *dunk* mutant embryos (He et al., 2016) and the two-hybrid interaction between Dunk and anillin prompted us to ask whether anillin colocalizes with myosin and/or regulates basal myosin array during early cellularization. It has been well documented that anillin colocalizes with myosin in the cytokinetic ring in many species and cell types during cytokinesis, including cellularization (Field et al., 2005; Piekny and Glotzer, 2008; Piekny and Maddox, 2010). However, the spatial distribution and function of anillin in cellularizing embryos prior to the formation of basal myosin rings are less well characterized. To determine whether anillin colocalizes with myosin during early cellularization, we performed live imaging analysis with embryos co-expressing GFP-tagged anillin (under the control of UAS-Gal4) (Silverman-Gavrila et al., 2008) and mCherry-tagged Sqh (Spaghetti squash, myosin regulatory light chain, under the control of the *sqh* promotor) (Martin et al., 2009). We found that anillin and myosin colocalized with each other at the basal contractile rings (Fig. 3; T = 30 – 45 min), as previously reported (Field et al., 2005). In addition, anillin and myosin also colocalized at the basal myosin array before the formation of myosin rings (Fig. 3; T = 5 – 25 min). A few minutes into cellularization, myosin started to form a basal array. The edges in the basal myosin array were first wide and fuzzy, and myosin signal appeared to be patchy (Fig. 3 and Fig. S1, T = 5 min). At this early stage, anillin was also enriched in the basal network and partially colocalized with myosin (Fig. 3 and Fig. S1, T = 5 min). As the edges of the basal array tightened over time, the colocalization between anillin and myosin became more prominent (Fig. 3, T = 10 – 25 min; Fig. S1, T = 15 and 25 min). Immunostaining of anillin and zipper, the myosin heavy chain, lead to similar observations (Fig. S2).

**Figure 3.**
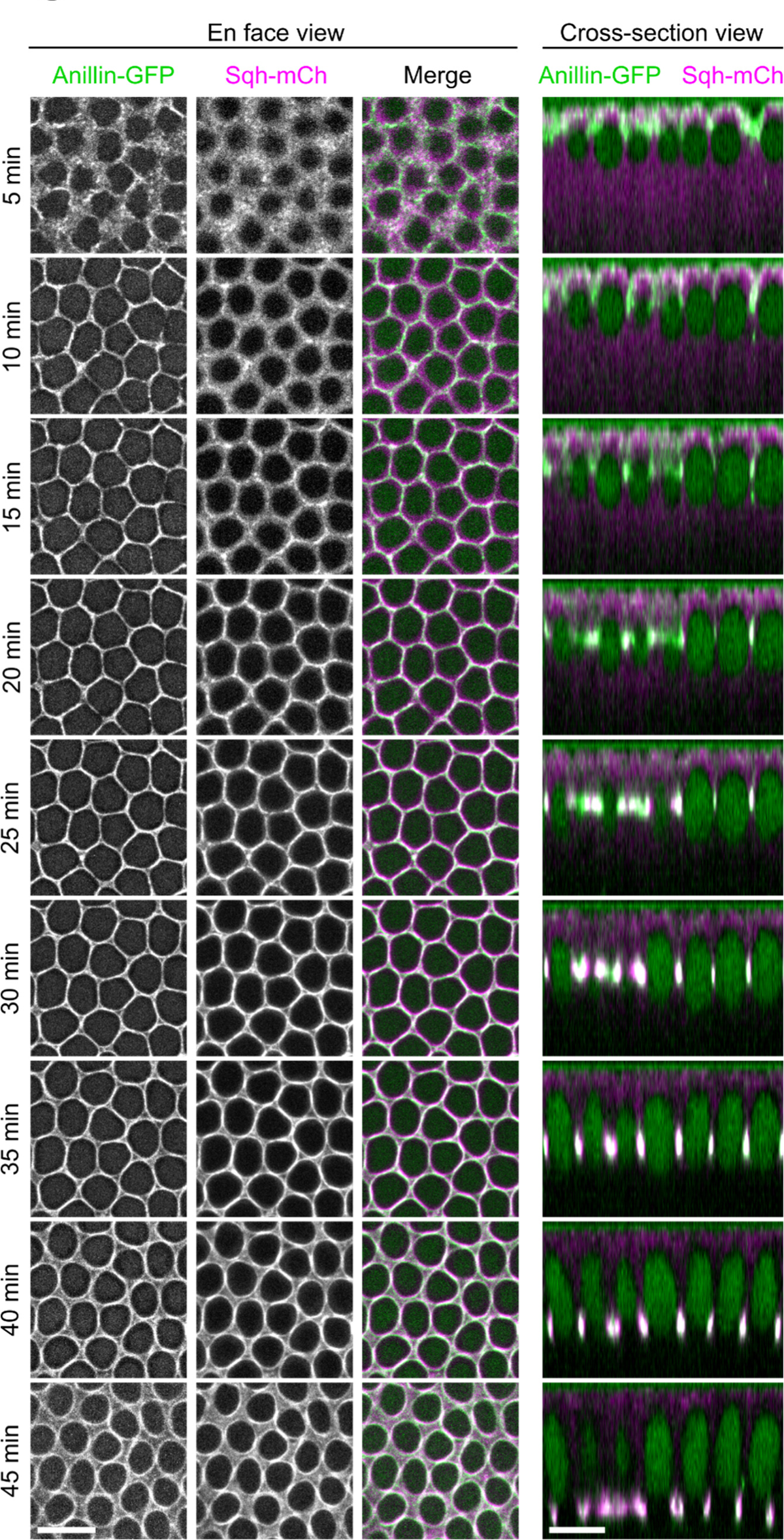
Localization of anillin and myosin during early and mid-cellularization. Live imaging of wild-type embryos expressing anillin-GFP and Sqh-mCherry. Left panels: en face view, maximum projections of 2 – 3 μm confocal sections covering the furrow canals; right panels: cross-section view generated by reslice of confocal z-stacks. The intensities of the en face views have been individually adjusted to better demonstrate the co-localization of the two proteins. Anillin and myosin partially colocalizes at the furrow canals during early cellularization (T = 5 min). The colocalization becomes more extensive after T = 10 min, and the two proteins remain colocalized throughout mid- to late cellularization as the basal array reorganizes into individual rings. Scale bars: 10 μm.

Interestingly, we found that the colocalization between myosin and anillin could be traced all the way back to the onset of cellularization (Fig 4A, T = 0 s, defined by the formation of daughter nuclei after the last syncytial nuclear division). At ∼ T = −2 min, myosin and anillin appeared and colocalized as punctum-like structures at the apical domain, enriched near nascent cleavage furrows (Fig 4A, colocalization: red arrow). Immediately after their first appearance, the puncta that were away from the nascent furrow moved towards the furrow (Fig. 4B, orange arrows), whereas those first appeared at the vicinity of the nascent furrow remained where they were (Fig. 4B, cyan arrows). Both behaviors contributed to the enrichment of myosin and anillin at the nascent furrows. Within an individual punctum, anillin and myosin appeared around the same time and remained colocalized throughout the cortical movement (Fig. 4B). The movement of myosin puncta towards the nascent furrow is consistent with the previous observation (He et al., 2016), and we further extended this observation to anillin by demonstrating the co-existence of myosin and anillin in the puncta. Taken together, our results demonstrate an extensive colocalization between anillin and myosin at the cellularization front both before and after the formation of basal myosin rings.

**Figure 4.**
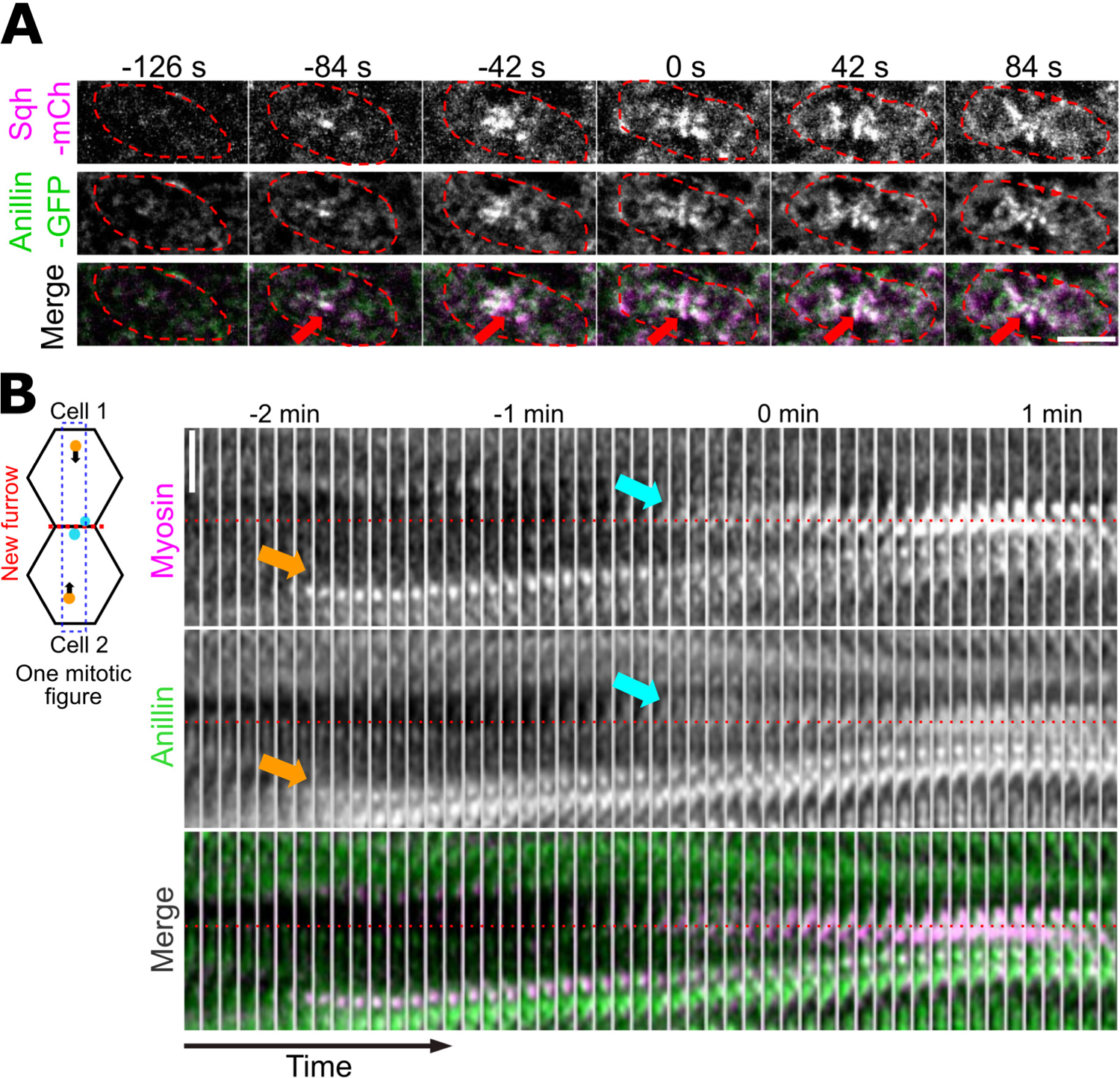
Anillin colocalizes and co-moves with cortical myosin puncta at the onset of cellularization. **(A)** Maximum projections of the apical z-slices (∼3 μm) of an embryo around the onset of cellularization (T = 0 s). Red arrows indicate the colocalization of anillin and myosin at the new furrow. Red dashed lines indicate the outline of the previous mitotic figure. Scale bar: 5 μm**. (B)** Left: A schema showing the apical anillin-myosin puncta in one mitotic figure. One mitotic figure contains two daughter cells. Some puncta (orange) first appear away from the nascent furrow (red dashed line) and move towards the new furrow (black arrows), whereas other puncta (cyan) first appeared at the nascent furrow. Blue box marks the region where the montage is generated. Right: Montage showing the cortical flow of anillin and myosin puncta towards the newly formed cleavage furrow at the beginning of cellularization. Orange and cyan arrows indicate puncta corresponding to the orange and cyan puncta in the schematics, respectively. Red dashed lines mark the position of the nascent cleavage furrow. Scale bar: 5 μm. Time scale: 4.2 sec per frame.

### Anillin regulates myosin localization at the furrow canals during early cellularization

The extensive colocalization of anillin with myosin during early cellularization prompted us to ask whether anillin regulates the formation and/or maintenance of the basal myosin array. To test this possibility, we imaged wildtype and *anillin* mutant embryos expressing GFP-tagged Sqh.

The *anillin* mutant embryos were derived from females that are trans-heterozygotes for two *anillin* mutant alleles (*anillin^PQ/RS^*, hereafter “*anillin* mutant”) (Schupbach and Wieschaus, 1989; Field et al., 2005). We first focused on the localization of myosin at the cellularization front after the initial recruitment of myosin to the nascent furrows (Fig. 5A). Compared to wildtype embryos, the spatial distribution of basal myosin was less uniform in *anillin* mutant embryos during the first 20 minutes of cellularization (Fig. 5A). Specifically, the myosin signal was aberrantly enriched at the vertices and largely depleted from the edges (Fig. 5A, 5 – 15 min, green and magenta arrows, respectively). As more myosin was recruited to the basal array over time, this “fragmented” basal myosin array phenotype became less prominent (Fig. 5A, 20 – 30 min). To further quantify the “fragmented” basal myosin phenotype, we segmented the basal array and measured the myosin intensities at the vertices and edges, as previously described (Fig. 5B; methods) (He et al., 2016). The average myosin intensity at the vertices is comparable between wildtype and *anillin* mutant embryos; however, there was less myosin at the edges in *anillin* mutant embryos compared to the wild type (Fig. 5C and E). As a result, the vertex/edge ratio of myosin intensity is higher in the *anillin* mutant embryos than that in the wildtype embryos (Fig. 5D and E). The defects in the spatial organization of basal myosin array, in particular the biased localization of myosin at the vertices and depletion of myosin from the vertices, closely resembles the reported myosin phenotype in *dunk* mutant embryos (He et al., 2016). This similarity is consistent with the notion that Dunk and anillin function in the same pathway to regulate cortical myosin organization during early cellularization.

**Figure 5.**
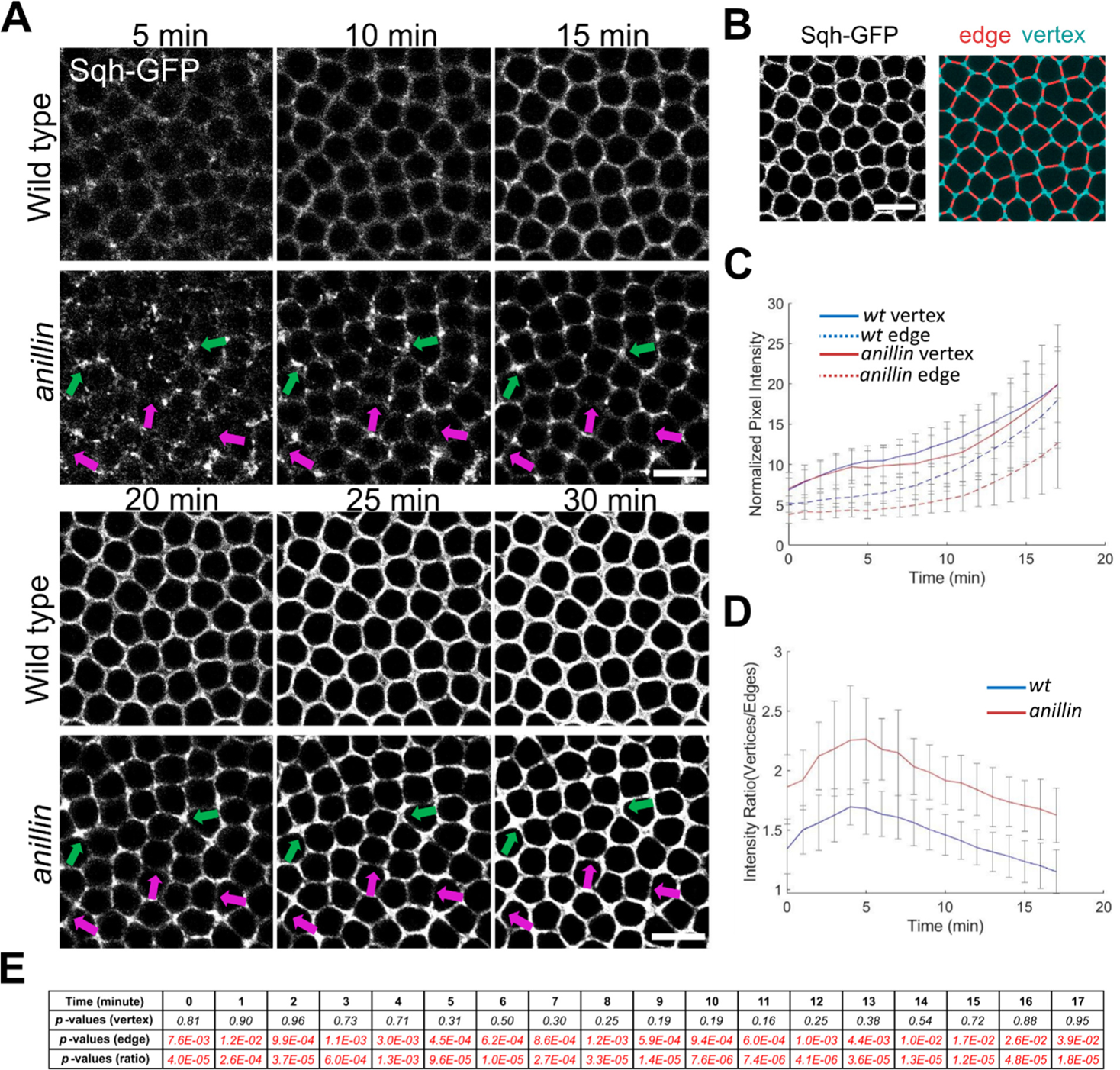
*anillin* mutant embryos show abnormal basal myosin localization during early cellularization. **(A)** Maximum projections of confocal sections showing Sqh-GFP at the furrow canals in wildtype and *anillin* maternal mutant embryos during the first 30 minutes of cellularization. Basal myosin distribution is abnormal in *anillin* mutant embryos. Specifically, during the first 20 min, myosin appears to be depleted from the edges (magenta arrows) and accumulated at some vertices (green arrows), closely resembling the previously reported myosin phenotypes in *dunk* mutant embryos (He et al., 2016). Scale bar: 10 μm. **(B)** Schematic diagram showing quantification of myosin intensity at vertices (cyan) and edges (red). Scale bar: 10 μm. (C) Quantification of vertex- and edge-myosin intensities at the furrow canals in wildtype (blue) and *anillin* mutant (red) embryos. **(D)** The ratio between vertex- and edge-myosin intensities. *Wild type*, *n* = 11 embryos; *anillin*, n = 12 embryos. Error bars: s.d. **(E)** Statistical comparison between wildtype and anillin embryos for mean intensity at the vertices, mean intensity at the edges, and the vertex/edge mean intensity ratios. Two-tailed unpaired student t-test was used. Red indicates statistically significant (*p* < 0.05). Black indicates no significant difference (*p* > 0.05).

Note that while the basal myosin phenotype in *anillin* mutants during early cellularization resembles that in *dunk* mutants, the myosin phenotype during late cellularization is different between the two genotypes. In *dunk* mutant embryos, basal myosin rings still form and contract to mediate basal closure of the nascent cells (He et al., 2016). In contrast, the proper myosin ring formation and subsequent basal closure are both defective in *anillin* mutant embryos (Field, 2005), suggesting that anillin has a separate function in late cellularization that is not dependent on Dunk.

### Anillin promotes the formation of cortical myosin puncta during the onset of cellularization

Next, we seek to elucidate how basal myosin phenotype develops during the first 5 – 10 minutes of cellularization. To test this, we examined wildtype and *anillin* mutant embryos expressing GFP-tagged Sqh at the transition between the syncytial and cellularization stages. We focused on the apical and subapical zones of the cell membrane from the time when myosin puncta first appeared until they moved away from the most part of the apical region. As previously described (He et al., 2016), in wildtype embryos, myosin first appeared at the apical domain of each mitotic figure when the nuclei started to form (T = −1 min), which was approximately 1 minute before the formation of the new furrow between the daughter nuclei (T = 0 min) (Fig. 6A). Meanwhile, myosin appeared at the base of the remnant pseudo-cleavage furrows (the old furrows, Fig. 6A, cyan arrowheads), which was retracting until ∼ T = 2 min when the new furrow reached the similar depth as the old furrows (Fig. 6A, orange box). The apical myosin puncta were initially quite sparse (T = −1 min) but the number increased rapidly and reached the peak at T = 1 – 2 min (Fig. 6A, cyan box; 6B). The puncta then quickly disappeared from the apical surface as the furrows invaginate. At T = 0 min, myosin puncta also appeared at the 1.5 – 3 μm zone and followed the same increase-decrease trend as the apical myosin puncta, but with a one-minute lag (Fig. 6A, green box). At T = 3 – 4 min, as myosin puncta have mostly disappeared from the 0 – 3 μm zone, myosin puncta became predominantly enriched at the 3 – 4.5 μm zone and uniformly decorated the leading edge of both old and new furrows (Fig. 6A, orange box). In the next few minutes, the edges of the basal myosin array tightened as the furrows continued to invaginate (Fig. 6A, magenta box).

**Figure 6.**
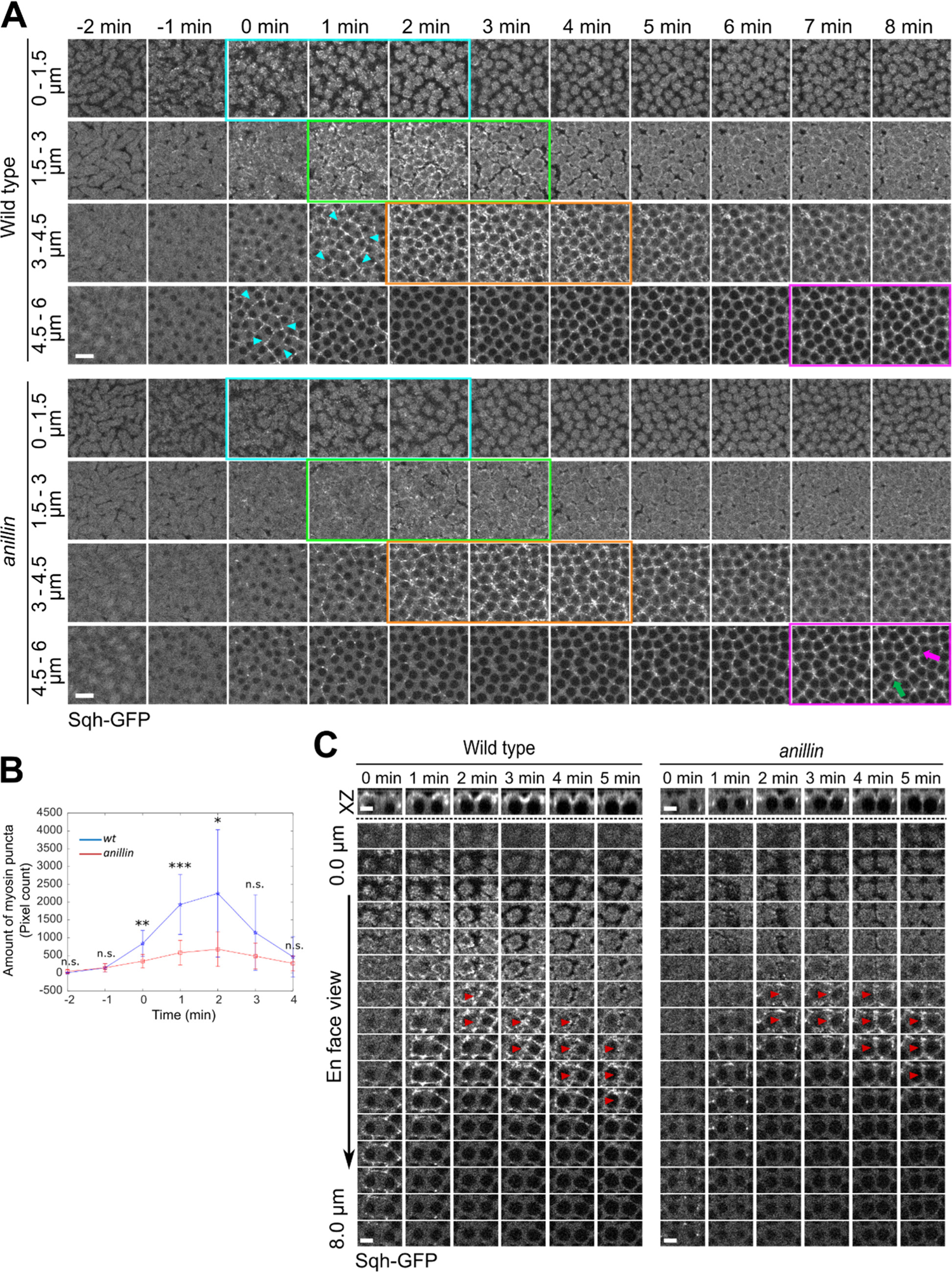
*anillin* mutant embryos show a reduced amount of apical myosin puncta and less stable basal myosin array at early cellularization. (A) Projections of confocal sections showing Sqh-GFP at the apical and subapical zones of the cell membrane (0 – 1.5 μm, 1.5 – 3 μm, 3 – 4.5 μm, and 4.5 – 6 μm) in wild-type and *anillin* mutant embryos over time. In wild-type embryos, myosin puncta first appear at the apical cortex (cyan box) and the tip of retracting pseudo-cleavage furrows (cyan arrowheads) around the onset of cellularization. The apical puncta rapidly disappear from the apical surface as the furrows invaginate (green box). At T = 2 – 4 min, as myosin puncta have mostly disappeared from the 0 – 3 μm zone, myosin puncta become predominantly enriched at the 3 – 4.5 μm zone and uniformly decorate the leading edge of both old and new furrows (orange box). In *anillin* mutant embryos, myosin puncta at the 0 – 3 μm region were greatly diminished compared to the wild-type, although the number of puncta showed a similar increase-decrease trend as the wild-type (cyan and green boxes). At T = 2 min, myosin started to be recruited to the cellularization front in *anillin* mutant at the 3 – 4.5 μm zone and was more or less uniformly distributed at the leading edge at this point (orange box). In the following several minutes, as the furrow continued to ingress, the myosin signal became progressively diminished at the edges (magenta arrow) and concentrated at the vertices (green arrow), resulting in the fragmented basal array phenotype (magenta boxes). Scale bar: 10 μm. **(B)** Quantification of myosin puncta at the apical cortex (0 – 2.5 μm) over time. The quantification measures the number of pixels of all myosin puncta in the view after segmenting the puncta by thresholding (methods). Error bars: s.d. A two-tailed unpaired student t-test was used for statistical comparison. *Wildtype*, *n* = 7 embryos; *anillin*, n = 10 embryos. *: *p* < 0.05; **: *p* < 0.01;*** *p* < 0.001: n.s., *p* > 0.05. **(C)** Examination of individual mitotic figures over time. Upper panel: Cross-section view of one mitotic figure. Lower panel: Montage of myosin signal over time at different depth. Red arrowhead: myosin was recruited to the base of new furrows in both *wildtype* and *anillin* mutant embryos but appeared to be less stable in *anillin* mutant embryos. Scale bar: 5 μm.

In *anillin* mutant embryos, we observed two major differences from the wildtype embryos. First, myosin puncta at the 0 – 3 μm region were greatly diminished compared to the wildtype, although the number of puncta showed a similar increase-decrease trend as in the wild type (Fig. 6A, cyan and green boxes; 6B). This result indicates that anillin facilitates the formation of apical myosin puncta at the onset of cellularization. Second, basal myosin appeared to be less stable in *anillin* mutant embryos compared to the wild type. At ∼ T = 2 min, myosin started to be recruited to the cellularization front at the 3 – 4.5 μm zone in *anillin* mutant embryos (Fig. 6A, orange box). Although the number of myosin puncta recruited to the leading edge appeared to be slightly fewer compared to the control embryos, these puncta were more or less uniformly distributed at the cellularization front. Examination of individual mitotic figures also shows that myosin was recruited to the base of both old and new furrows (Fig. 6C, red arrowheads indicate myosin in the new furrow). However, in the following several minutes, as the furrows continued to ingress (i.e., basal myosin moved deeper into the embryo), myosin signal became progressively diminished at the edges and concentrated at the vertices, resulting in the fragmented basal myosin array phenotype described in Figure 5 (Fig. 6A, magenta box, magenta and green arrows show depletion and enrichment of the myosin signal, respectively). These results suggest that anillin have two roles during the flow phase of myosin recruitment. First, anillin is required for the formation of apical myosin puncta at the onset of cellularization. The flow of these myosin puncta to the nascent furrows contributes to initial myosin recruitment to the cellularization front. Second, anillin facilitates the retention of myosin at the base of the ingressing furrows, which is important for the formation of an interconnected basal myosin array and the subsequent assembly of myosin rings.

### Genetic interaction between *anillin* and *dunk*

To further test whether Dunk and anillin function together to regulate basal myosin during early cellularization, we examined potential genetic interactions between the two genes. Previously identified *anillin* and *dunk* mutants are all characterized as recessive alleles. No obvious phenotypes have been reported for *anillin* or *dunk* heterozygous mutants (Field et al., 2005; He et al., 2016). If anillin and Dunk function in the same pathway regulating basal myosin network during cellularization, simultaneously reducing the amount of both proteins by double heterozygosity might impair the pathway and result in detectable lesions. To test this, we generated *anillin dunk* double heterozygous mutant and asked whether it shows any synthetic phenotypes during cellularization. In the analysis described below, the *anillin* heterozygous (*ani*/*+*) refers to the maternal genotype, whereas the *dunk* heterozygous (*dunk*/*+*) refers to the zygotic genotype (Methods). We first acquired snapshots of basal myosin in embryos at mid-cellularization when myosin rings have formed in wildtype embryos. We found that mutating one copy of *anillin* or *dunk* did not have a significant impact on basal myosin rings (Fig. 7A).

**Figure 7.**
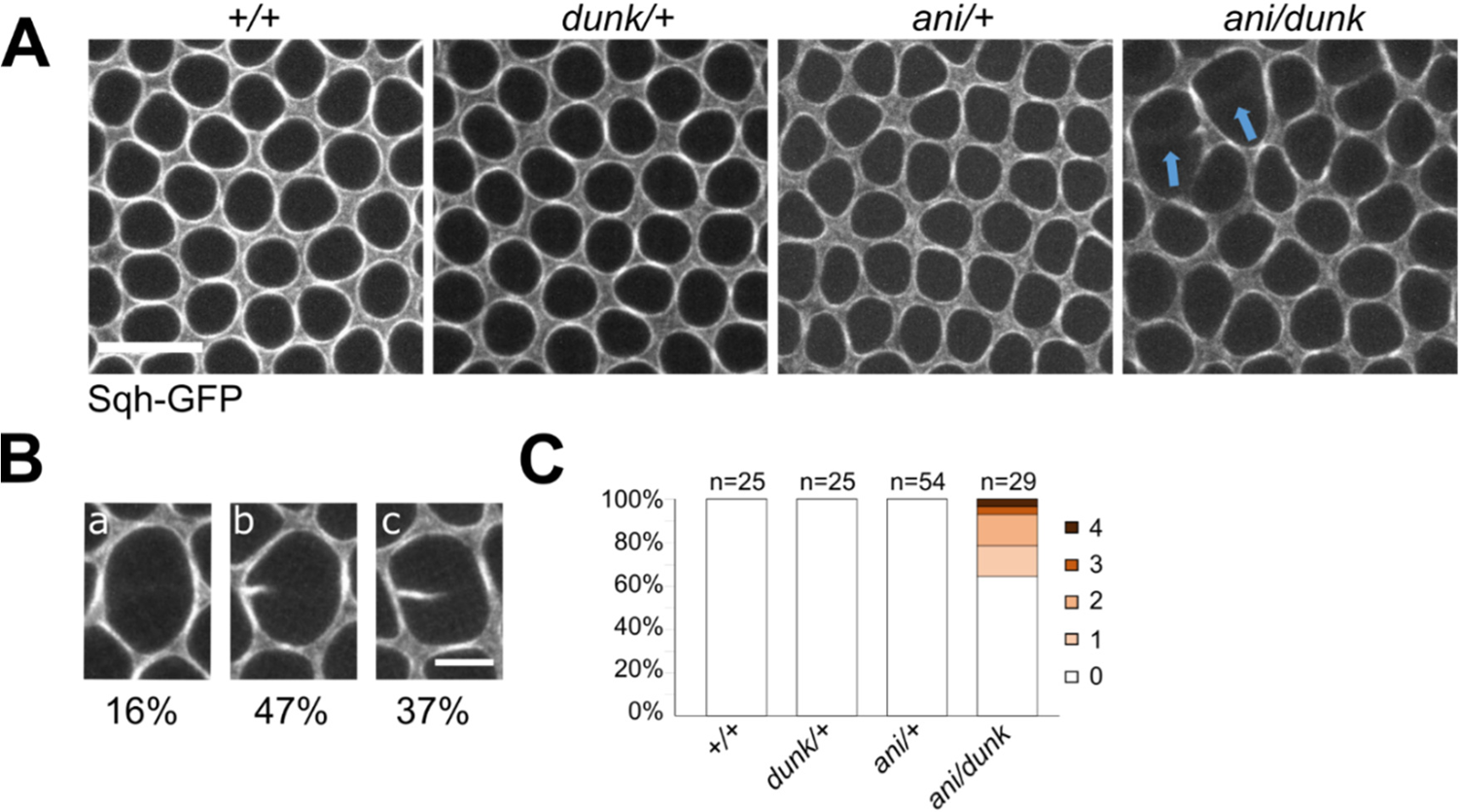
Genetic interaction between *dunk* and *anillin*. **(A)** Basal myosin organization in wild type (*+/+*), *dunk* heterozygous mutant (*dunk/+*), *anillin* heterozygous mutant (*ani/+*) and *anillin dunk* double heterozygous mutant (*ani/dunk*). ‘*anillin*’ here refers to the maternal genotype and ‘*dunk*’ here refers to the zygotic genotype. Shown are representative images of basal myosin rings in mid- to late cellularization. The “broken-myosin-edge” phenotype, which refers to the loss of basal myosin signal from at least half of an edge between a pair of neighboring cells, was only observed in the double heterozygous mutant (blue arrows). Scale bar: 10 μm. **(B)** Enlarged view of different types of “broken-myosin-edge” phenotype observed in the double heterozygous mutant embryos. a, b and c show loss of myosin signal from the entire edge, 66.7% – 100% of the edge and 50% – 66.7% of the edge, respectively. The numbers below the images show the frequency of each type. **(C)** The fraction of embryos showing 1 – 4 broken-myosin-edges in the imaged region (∼5% of the entire embryo surface). The “broken-myosin-edge” phenotype only exits in double heterozygous mutant embryos. The numbers of embryos analyzed for each genotype are shown in the figure.

Interestingly, in about a third of the *anillin*/*dunk* double heterozygous embryos (10 out of 29 embryos; the imaged region covered ∼5% of the total embryo surface), when basal myosin first reorganized into individual rings, we observed “broken” myosin edges between one or more pairs of neighboring cells (Fig. 7A, arrows). In some cases, the entire edge was missing (Fig. 7B, “a”). In other cases, myosin was missing from a part of the edge (Fig. 6D, “b” and “c”: missing from 66.7% − 100% and 50% − 66.7% of the edge, respectively). This “broken-myosin-edge” phenotype was only observed in the double heterozygous mutant, but not in the wild type or either of the single heterozygous mutants (Fig. 7C). Therefore, simultaneously reducing the amount of Dunk and anillin to the level of heterozygous mutant generates a synthetic effect on basal myosin organization during cellularization.

We next wondered whether the synthetic effect on myosin organization can be detected at an earlier stage before the basal myosin network reorganized into discrete contractile rings. To this end, we acquired movies that covered early cellularization stages (T = 0 – 10 min). In both single and double heterozygous mutant embryos, the basal myosin array formed at about the same time as in wildtype embryos, but subtle differences in the arrangement of basal myosin array could be detected between genotypes (Fig. S3A). While the localization of myosin at the basal array in *dunk*/+ embryos was nearly identical to that in wildtype embryos, the localization of myosin is more heterogeneous in *ani*/*+* and *anillin*/*dunk* embryos, with a biased enrichment at the vertices and depletion from the edges (Fig. S3A, green and magenta arrows, respectively). The biased distribution of myosin was more prominent in *anillin*/*dunk* embryos than in *anillin*/+ embryos, although the embryo-to-embryo variations were relatively large in both populations (Fig. S3A, “mild” and “severe”) compared to wildtype and *dunk*/*+* groups. Quantification of the vertex/edge ratio of myosin intensity at T = 10 min for each genotype confirmed our observations, showing that *ani/+* has a higher vertex/edge intensity ratio than the wild type and *dunk/+*, whereas *anillin dunk* double heterozygous mutant had the highest vertex/edge intensity ratio (Fig. S3B). Together, the observed additive effects of *ani/+* and *dunk/+* on basal myosin organization support the notion that Dunk and anillin function together to regulate basal myosin during cellularization.

### *dunk bnk* double mutants show similar myosin phenotype during cellularization as *anillin bnk* double mutants

During cellularization, the reorganization of the basal myosin network into individual contractile myosin rings is regulated by *bottleneck* (*bnk*) (Schejter and Wieschaus, 1993). In *bnk* mutant embryos, the basal actomyosin array reorganizes into contractile rings prematurely and displays hyper-contractility (Fig. 8A’, B’) (Schejter and Wieschaus, 1993). A previous study shows that when *anillin* and *bnk* are both mutated, the phenotypes of each single mutant are co-expressed: in the absence of anillin, the structural organization of basal myosin is impaired, whereas a further increase in actomyosin contractility caused by lack of Bnk results in rupturing of the basal network in cellularization (Thomas and Wieschaus, 2004). We reasoned that if Dunk and anillin function in the same pathway in early cellularization, combining mutations in *dunk* and *bnk* would generate a similar phenotype. Indeed, we found that unlike the *dunk* mutant embryos where the basal array appeared intact without obvious ruptures (comparing Fig. 8C’ and 8A’), the basal network in *bnk dunk* mutant embryos was severely disrupted, yielding large holes at the furrow canals that encloses multiple nuclei (Fig. 8D’, magenta arrows). These holes were usually circular, suggesting that the ruptured basal array was still under tension. In addition, at regions where rupture did not occur, basal myosin displayed hyper-constricted phenotype as in *bnk* mutant embryos (comparing Fig. 8D’ with Fig. 8B’, cyan arrowheads). These observations demonstrate that similar to *bnk* and *anillin*, the *bnk* and *dunk* phenotypes are also co-expressed in the double mutant embryos. The *bnk dunk* double mutant phenotype is in sharp contrast to that of the *bnk src64* double mutant, in which the *bnk* phenotype is suppressed by removal of Src64, a positive regulator of actomyosin contractility (Thomas and Wieschaus, 2004). These results are consistent with the notion that Dunk does not function to activate myosin contractility, but rather functions to maintain the structural integrity of the basal actomyosin structures (He et al., 2016). The similarity in the basal myosin phenotype between *dunk bnk* and *anillin bnk* double mutants provides further support that Dunk and anillin function in a common pathway that regulates basal myosin organization during early cellularization.

**Figure 8.**
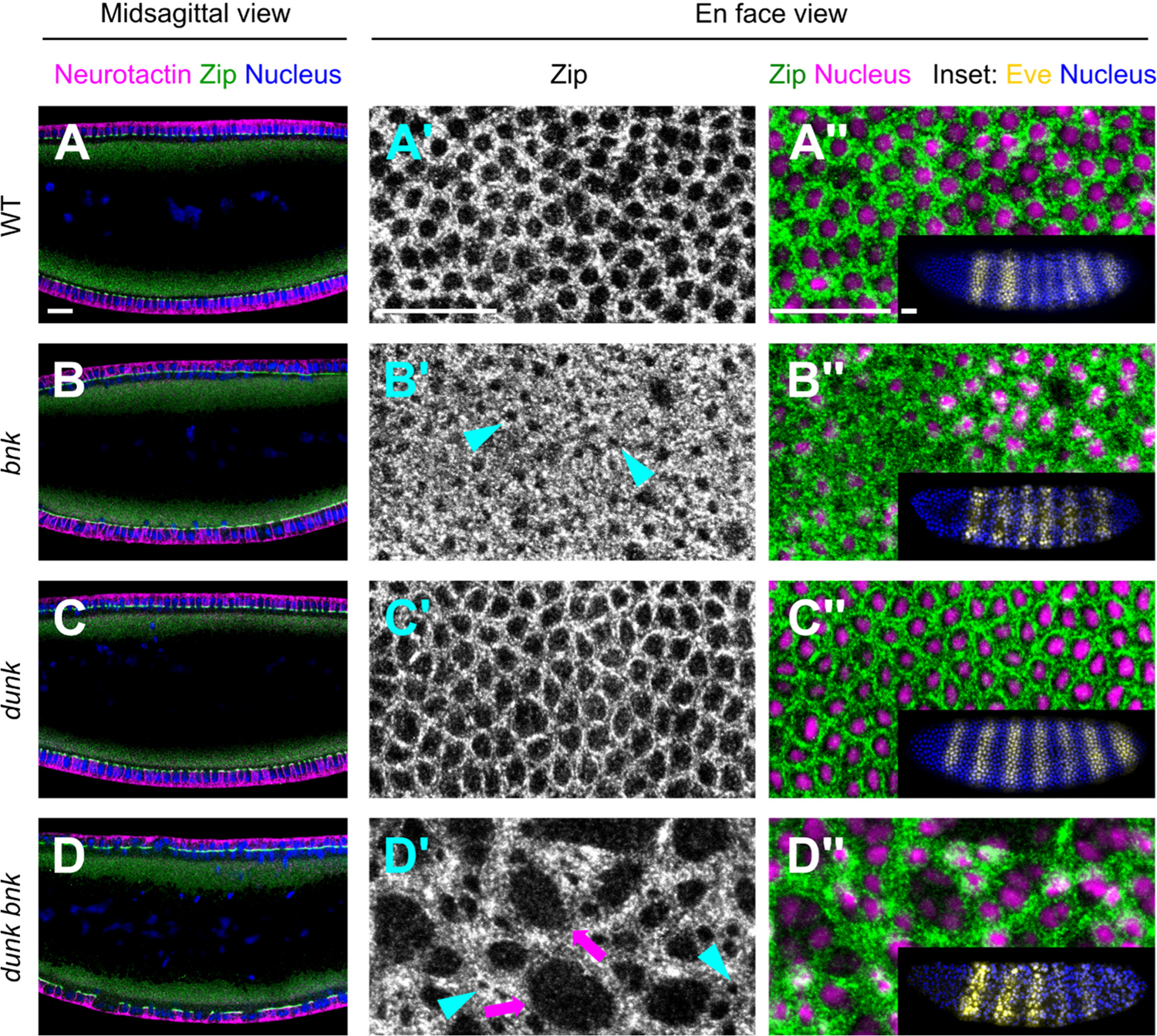
Increasing basal actomyosin contractility exacerbates the myosin phenotype in *dunk^1^* mutant embryos. **(A)** wildtype embryo. **(B)** *bnk* mutant embryo. **(C)** *dunk^1^* mutant embryo. **(D)** *bnk dunk^1^* double mutant embryo. Left column: Cross-sections of embryos showing staining for membrane (Neurotactin), myosin (Zipper) and nucleus. Middle and Right columns: Projections of confocal sections showing the en face view of the furrow canals as marked by myosin (Zipper). The phenotypes of *dunk* and *bnk* are co-expressed. The disruption of *dunk* in *bnk* mutant leads to a combined phenotype of premature ring contractions (*bnk*, cyan arrowheads) and reduced integrity of the basal actomyosin network (*dunk*), causing the tearing of the network (magenta arrows). Inset on the right: Immunostaining of Even skipped (Eve), which was used to identify *bnk* mutant embryos (Methods). *Df(3R)^tll-e^*, the deficiency used to generate *bnk* mutant embryos, contains a deletion that covers both *bnk* and the terminal gap gene *tll* (Schejter and Wieschaus, 1993). Tll regulates the expression of pair-rule genes in a stereotyped manner (Frasch and Levine, 1987). *tll* mutant embryos show 6 Eve strips instead of 7 as in the wild-type embryos. All scale bars: 20 µm.

## Discussion

Proper recruitment of myosin to the cleavage furrow is a critical step in animal cytokinesis (Green et al., 2012; Normand and King, 2010; Piekny and Maddox, 2010). *Drosophila* cellularization features the synchronized formation of several thousands of cleavage furrows from the surface of the embryo. This process is accompanied by rapid, simultaneous recruitment of myosin to the entire array of nascent cleavage furrows, which makes cellularization an advantageous system to study the mechanism of myosin recruitment and organization during cytokinesis (Royou et al., 2004; He et al., 2016). Previous identification of *dunk*, a cellularization-specific gene that regulates the initial recruitment of myosin to the basal array, provides a useful molecular handle to investigate the mechanism of this process (He et al., 2016). In this study, we present evidence that Dunk functions together with anillin, a well-conserved scaffolding protein in cytokinesis, to regulate basal myosin during early cellularization (Field and Alberts, 1995; Field et al., 2005; Piekny and Glotzer, 2008). Through a genome-wide yeast two-hybrid screen, we identified anillin as the sole binding partner for Dunk. Further domain analysis demonstrates that Dunk binds to the C-terminal region of anillin, and this binding requires both the AH and PH domains. We further present evidence supporting a functional interaction of Dunk with anillin in regulating basal myosin. Loss of Dunk results in mislocalization of anillin and septin at the basal array and causes defects in the formation of basal anillin and septin rings. Mutant embryos that are heterozygous for both *dunk* and *anillin* loss of function mutations display synthetic phenotypes in basal myosin organization. Furthermore, *dunk* and *anillin* mutants show identical synthetic phenotypes when combined with the mutation in *bottleneck* that enhances contractility of the basal actomyosin network. Finally, we found that anillin colocalizes with myosin at the cellularization front during early cellularization and regulates myosin recruitment and retention at the basal array. This previously unappreciated function of anillin during early cellularization is similar to the reported function of Dunk (He et al., 2016). Together, these results led us to propose that Dunk regulates myosin at the furrow canals by interacting with anillin and controlling anillin’s localization and/or activity.

Although the function of anillin in regulating actomyosin ring during cytokinesis has been well demonstrated (Field et al., 2005; Piekny and Glotzer, 2008; Piekny and Maddox, 2010), our understanding of its function prior to ring formation remains limited. Our quantitative analysis of Sqh-GFP shows that in addition to the well-characterized role of anillin in regulating myosin ring assembly, anillin also regulates myosin recruitment and maintenance at the furrow canals prior to myosin ring formation. In the absence of functional anillin, less myosin is recruited to the apical cortex at the onset of cellularization. Furthermore, myosin recruited to the basal array becomes less stable, causing aberrant spatial distribution of myosin across the leading edge of the furrows. This function of anillin is consistent with the observation that anillin not only colocalizes with the basal myosin rings during mid-late cellularization but also colocalizes with the basal myosin array during early cellularization. Most interestingly, we found that when myosin was first recruited to the apical cell cortex at the onset of cellularization, anillin colocalizes with myosin as discrete cortical puncta and co-move with myosin towards the nascent cleavage furrow. The cortical flow of myosin towards the prospective cleavage furrow has been observed in various cytokinesis contexts (DeBiasio et al., 1996; He et al., 2016; Uehara et al., 2010; Yumura et al., 2008). It would be of interest to test whether anillin is also involved in cortical myosin flow during other cytokinesis processes and whether other cytokinetic proteins are co-transported with the flow.

It remains to be addressed how anillin promotes myosin recruitment or cortical retention during early cellularization, and how Dunk regulates this process. Anillin may facilitate the cortical localization of myosin through direct interaction. Alternatively, anillin may help activate myosin through its interaction with RhoA. A recent study suggests that anillin promotes RhoA activation by increasing the membrane residence time of RhoA (Budnar et al., 2019). Since Dunk is predicted to bind to the conserved AH and PH domains of anillin, Dunk may regulate anillin in a synergistic manner with Rho1 and PI(4,5)P_2_. Interestingly, previous work has shown that although Dunk promotes the cortical retention of myosin during early cellularization, loss of Dunk does not affect the rate of myosin turnover at the cell cortex (He et al., 2016). This leads to the hypothesis that Dunk may regulate myosin retention by controlling the available myosin-binding sites at the cell cortex, rather than controlling the dynamic interaction of myosin with the binding sites per se. The observation that the accumulation of cortical myosin puncta at the onset of cellularization reduces in *anillin* mutant embryos raises the possibility that anillin at the cell cortex may serve as cortical myosin-binding sites or function to stabilize such sites during the early phase of cellularization.

Notably, the defect in myosin accumulation seen in the *anillin* and *dunk* mutant embryos became less prominent after the first 20 – 25 minutes of cellularization (Field et al., 2005; He et al., 2016), suggesting that certain anillin- and Dunk-independent mechanisms for myosin activation/recruitment (e.g. Slam-dependent direct recruitment of myosin) is in charge after the initial phase of cellularization. In addition, despite the similarity of the myosin phenotype in *dunk* and *anillin* mutant embryos during early cellularization, the *anillin* mutant, but not the *dunk* mutant, blocks the rearrangement of basal myosin into individual rings during mid cellularization (Field et al., 2005; He et al., 2016). Therefore, while the early function of anillin in regulating myosin enrichment at the furrow canals is shared with Dunk, the late function of anillin in regulating myosin ring formation is independent of Dunk. The unique, temporally separable mechanisms for myosin recruitment and rearrangement during cellularization (He et al., 2016; Xue and Sokac, 2016) makes it an attractive system to reveal the stage-specific function and regulation of anillin during cytokinesis.

Like other cellularization-specific proteins, Dunk is not conserved in other model organisms. Sequence homologs of Dunk are only found in flies and related insects. This leads to the interesting question of why Dunk is required for regulating anillin in cellularization but not in conventional cytokinesis, including post-cellularization cytokinesis in flies. Despite all the similarities, cellularization is different from typical cytokinesis in a number of ways. One important difference is the cell cycle stage when the cleavage furrows are formed. Cellularization happens during the interphase of the cell cycle (Foe and Alberts, 1983), while typical cytokinesis happens during late anaphase to telophase (Green et al., 2012). In typical cytokinesis, anillin localizes to the nucleus during interphase and is recruited to the cell membrane during mitosis.

When cellularization starts, however, anillin is immediately recruited to the nascent cleavage furrows, in contrast to a slower enrichment in the nucleus (Field et al., 2005). An intriguing hypothesis is that the function of Dunk is required to adapt the behavior of anillin during the normal cell cycle in order to support cytokinesis during interphase. Future research will be needed to test this hypothesis.

## Materials and Methods

### Yeast two-hybrid assay

Full-length *dunk* was cloned into pGBKT-7 plasmid (a vector containing DNA-binding domain (BD) of GAL4) and transformed into Y2H Gold yeast cell as the bait for all the yeast two-hybrid experiments. The expression of Dunk protein was confirmed by Western blot using c-Myc Monoclonal Antibody (Takara, Cat# 631206). Then, the Y2HGold[pGBKT7-Dunk] was tested for autoactivation and toxicity. The results are described below. First, Y2Hgold[pGBKT7-Dunk] yeast colonies on the SDO/X plates were in white color, indicating that Dunk-BD alone cannot activate the transcription of the reporter gene *MEL1*, whose product can hydrolyze colorless X-alpha-Gal into a product with blue color. Second, Y2HGold[pGBKT7-Dunk] could not survive on SDO/X/A plates, indicating that Dunk-BD alone cannot activate the transcription of the reporter gene *AUR1-C*, which is required for survival on the plates with Aureobasidin A. Finally, the growth of Y2HGold[pGBKT7-Dunk] cell was comparable to Y2HGold[pGBKT7], indicating the Dunk protein is not toxic when expressed in yeast.

To perform the screen for Dunk-binding proteins, a full-length Dunk was used to screen the Mate & Plate™ Library - Universal *Drosophila* (Normalized) (Takara, 630485). This genome-wide *Drosophila* cDNA library is prepared from equal quantities of poly-A^+^ RNA isolated from embryo, larva and adult stage *Drosophila*, covering most of the expressed genes in *Drosophila*. The gene representation has been equalized by reducing the copy number of abundant cDNAs which in turn increased the occurrence of low copy number transcripts (Zhulidov et al., 2004).

This *Drosophila* cDNA library was cloned into pGADT-7 plasmid (a vector containing activation domain (AD) of GAL4) and transformed into yeast strain Y187. The screen was performed followed by the Matchmaker Gold Yeast Two-Hybrid System User Manual (Takara, 071519). One large Y2HGold[pGBKT7-Dunk] yeast colony was picked from a fresh SD/-Trp plate and inoculated into 50 ml of SD/-Trp liquid media. The culture was incubated and shook (250–270 rpm) at 30°C until the OD_600_ reaches 0.8 (16–20 hr). Y2HGold[pGBKT7-Dunk] yeast cells were pelleted by centrifuging at 1000 g for 5 min. The pellet was resuspended to a cell density of >1 x 10^8^ cells per ml in SD/–Trp (4–5 ml). The cell density of Y2HGold[pGBKT7-Dunk] was checked by the hemocytometer under the phase-contrast microscope. After checking the quality of the library strain by titering on SD/-Leu plates (1 ml library should contain more than 2 x 10^7^ cells), 1 ml of the library strain was combined with the Y2HGold[pGBKT7-Dunk] strain in a sterile 2 L flask with 45 ml 2×YPDA liquid medium (with 50 µg/ml kanamycin). The culture was incubated and shook at 30°C for 20 hr at 50 rpm. The formation of the zygotes was checked under the phase-contrast microscope after 20 hr, indicating the mating between the bait strain and the library strain. Then cells were pelleted by centrifuging at 1000 g for 10 min. Resuspended the cell pellet with 50 ml 0.5× YPDA (with 50 µg/ml kanamycin) and centrifuged to pellet the cells again. The cell pellet was resuspended again with 10 ml 0.5× YPDA (with 50 µg/ml kanamycin). The total volume of the media and the cell are measured (TV). 200 µl of the culture was plated on per 150 mm DDO/X/A plate. The plates were incubated at 30°C for 3-5 days. At the same time, spread 100 µl of 1/10, 1/100, 1/1,000, and 1/10,000 dilutions of the mated culture on DDO 100 mm agar plates and incubate at 30°C for 3–5 days. The numbers of screened colonies were calculated by the numbers of diploids that grew on the DDO plates (Num), resuspension volume (TV), plating volume (PV) and the dilution factor (D). Numbers of screened colonies = Num × TV / (PV × D). No less than 1 million diploids should be screened.

After 3-5 days, positive colonies (blue colonies) from the DDO/X/A plates were picked, and the plasmids purified from these positive clones were sequenced using the sequencing primers for pGADT-7 plasmid. We have repeated this genome-wide screen three times and in total 7 positive colonies were identified and sequenced. All the prey plasmids in those positive colonies contained the C-terminal of *Drosophila* anillin (630 – 1212 a.a.).

For the control, pGBKT7-53 (encodes the Gal4 DNA-BD fused with murine p53) and pGBKT7-Lam (which encodes the Gal4 BD fused with lamin) constructs were transformed into Y2H Gold yeast cell. pGADT7-T (encodes the Gal4 AD fused with SV40 large T-antigen) vector was transformed into Y187 yeast cell. Since p53 and large T-antigen are known to interact in a yeast two-hybrid assay (Li & Fields, 1993; Iwabuchi et al. 1993), mating Y2HGold [pGBKT7-53] with Y187 [pGADT7-T] was treated as the positive control for the screen. Since lamin does not interact with T-antigen in a yeast two-hybrid assay, Y2HGold [pGBKT7-lam] with Y187 [pGADT7-T] was treated as the negative control for the screen.

For identifying Dunk binding domain on anillin, Scraps truncations ani_CT (592-1212 a.a.), ani_PH (1000-1212 a.a.), ani_CTΔPH (592-1000 a.a.), ani_ND (592-820 a.a.) and ani_AHPH (820-1212 a.a.) were cloned into pGADT-7 plasmid and transformed into yeast strain Y187 as preys. The truncations were designed based on Scraps isoform A since this isoform has the same C-terminal sequence as the prey strain we found in the genome-wide screen. The different anillin truncations were tested for binding to full-length Dunk using yeast two-hybrid. After the mating, diploids were spotted on the SD/–Leu/–Trp/X-alpha-Gal/AbA agar plates. The appearance and the survival of the colonies indicate the interaction between the prey and the bait proteins, and the intensity of the blue color of the colonies indicates the strength of the interaction.

### Fly stocks and genetics

Fly lines containing the following fluorescent markers were used: *UAS-GFP-anillin* (*UAS-GFP-scraps*) (Silverman-Gavrila et al., 2008), *Sqh-mCherry* (Martin et al., 2009) and *Sqh-GFP* (Royou et al., 2002). The *Maternal-Tubulin-Gal4* line 67.15 (“*67*” and “*15*” refer to *Maternal-Tubulin-Gal4* on chromosome II and III, respectively) (Hunter and Wieschaus, 2000) was used to drive the expression of GFP-anillin.

*OreR* embryos were used as a control for immunostaining experiments unless stated otherwise. The *dunk^1^* P-element insertion mutant line, *P{SUPor-P}CG42748^KG09309^* (Bellen et al., 2004), was obtained from Bloomington *Drosophila* Stock Center.

For examining the colocalization of myosin and anillin, UAS-GFP-anillin (III) females were crossed to males from the Maternal-Tubulin-Gal4-15 sqh-mCherry (III) to generate UAS-GFP-anillin (III)/ Maternal-Tubulin-Gal4-15 sqh-mCherry (III) flies. Embryos derived from these flies were used to examine the localization of anillin and myosin by live imaging.

For examining myosin localization in *anillin* mutant embryos, *scra^PQ^/Cyo*; *Sqh-GFP* females were crossed to *scra^RS^/Cyo*; *Sqh-GFP* males to generate *scra^PQ^/scra^RS^*; *Sqh-GFP* transheterozygous flies (Schupbach and Wieschaus, 1989). Embryos derived from these flies were used to examine myosin localization by live imaging. *scra^PQ^* and *scra^RS^* are maternal-effect alleles containing point mutations in the PH domain (Field et al., 2005).

The following crosses were made to generate (1) *ani*/*+*, (2) *dunk^1^*/+, and (3) *ani*/*dunk^1^* heterozygous mutant embryos. Note that *scra^PQ^* and *Sqh-GFP* are the maternal genotype and *dunk* is the zygotic genotype. (1) Females from *scra^PQ^*/*Cyo(II)*; *Sqh-GFP (III)* were crossed to *OreR* males to generate *scra^PQ^ /+ (II)*; *Sqh-GFP (III)* embryos. In these embryos, the amount of maternally deposited, functional anillin is expected to be 50% of those in the wildtype embryos. (2) Females from *dunk^1^(II)*; *Sqh-GFP (III)* were crossed to *OreR* males to generate *dunk^1^ /+ (II)*; *Sqh-GFP (III)* embryos. In these embryos, the amount of zygotically expressed Dunk is expected to be 50% of those in the wildtype embryos. (3) Females from *scra^PQ^*/*Cyo(II)*; *Sqh-GFP (III)* were crossed to *dunk^1^(II)*; *Sqh-GFP (III)* males to generate *dunk^1^ / scra^PQ^ (II)*; *Sqh-GFP (III)* embryos. In these embryos, the amount of maternally deposited, functional anillin and the amount of zygotically expressed Dunk are expected to be 50% of those in the wildtype embryos. The localization of Sqh-GFP is examined in these three types of embryos and the wildtype embryos.

The *dunk bnk* double mutant embryos were generated from the *dunk^1^*; *Df(3R)^tll-e^*/TM3 flies. A quarter of the embryos are expected to be *dunk^1^*/*dunk^1^*; *Df(3R)^tll-e^*/*Df(3R)^tll-e^*. *Df(3R)^tll-e^* contains a deletion that covers both *bnk* and the terminal gap gene *tll* (Schejter and Wieschaus, 1993). Tll regulates the expression of pair-rule genes in a stereotyped manner (Frasch and Levine, 1987). Thus, embryos homozygous for this deficiency can be identified by the immunostaining of the products of pair-rule genes, such as Even skipped (Eve).

### Embryo fixation and antibody staining

Antibody staining against myosin (Zipper) and neurotactin (Nrt) was performed on heat-fixed embryos. Antibody staining against Scraps and Peanut (Pnut) was performed on formaldehyde-fixed embryos. The vitelline membrane was removed by shaking in heptane and methanol after fixation. Embryos were blocked with 10% BSA in PBS and 0.1% Tween 20, and incubated with primary antibodies in PBT (PBS/0.1% BSA/0.1% Tween 20) overnight at 4°C with the following dilutions: rabbit anti-Zipper 1:100; monoclonal mouse anti-Nrt 1:10 (BP 106, Developmental Studies Hybridoma Bank); rabbit anti-Scraps 1:1000 (gift of C. Field); monoclonal mouse anti-Pnut 1:10 (4C9H4, Developmental Studies Hybridoma Bank). Secondary antibodies coupled to Alexa488, Alexa561, and Alexa647 were used at 1:500 (Invitrogen). Embryos were mounted in Aqua Poly Mount (Polysciences) for confocal imaging. Confocal images were collected on a Leica SP5 confocal microscope with a 63×/1.3 NA glycerine-immersion objective lens and a pinhole setting of 1 airy unit.

### Live imaging

To prepare embryos for live imaging, manually staged embryos expressing anillin-GFP/ sqh-mCherry or sqh-GFP were collected at room temperature (22-25°C) on agar plates, dechorionated in 50% bleach for 1-2 min, rinsed thoroughly with water, and transferred on a 35 mm MatTek glass-bottom dish (MatTek Corporation). Distilled water was then added to the dish well to completely cover the embryos. All live imaging was performed at room temperature on a Nikon inverted spinning disk confocal microscope equipped with Andor W1 dual camera, dual spinning disk module. A CFI Plan Apo Lambda 60×/1.40 WD 0.13 mm Oil Objective Lens objective was used for imaging. GFP- and mCherry-tagged proteins were imaged with a 488-nm laser and a 561-nm laser, respectively.

### Image analysis and quantification

To compare myosin fluorescence intensity at the edges and vertices (Fig. 5), the Sqh-GFP movies were analyzed using MATLAB (Image Processing Toolbox, The MathWorks; Natick, MA) as follows. First, maximum intensity projections were generated from raw images from 7 adjacent confocal slices (0.5 µm z-step, ∼ 3 µm thick) that cover the invagination front. Second, the projected images were subject to image background subtraction. Third, to define signals that belong to edges versus vertices, the basal outline of the cells (as marked by Sqh-GFP) were segmented using a MATLAB-based software package Embryo Development Geometry Explorer (EDGE) (Gelbart et al., 2012). In EDGE, the outlines of individual cells are represented by polygons and tracked over time. Along each polygon, we define points less than 1.2 µm away from the nearest vertex as “vertex”, whereas points more than 1.2 µm away from the nearest vertex as “edge”. Mean intensity was integrated at vertices and edges along with the corresponding line segments with a width of 0.3 µm. The median pixel intensity of the image is used as a proxy for the intensity of the cytoplasmic Sqh-GFP signal and is subtracted from the mean vertex and edge intensities. Finally, the intensity was normalized between embryos according to the median pixel intensity of the image.

To compare myosin puncta fluorescence intensity in wildtype and *anillin* mutant (Fig. 6), the Sqh-GFP movies were analyzed using MATLAB as follows. The maximum intensity projections were generated from raw images from 5 adjacent confocal slices (0.5 µm z-step, ∼ 2.5 µm thick) that cover the most apical cortex. The background subtraction and intensity normalization are performed in the same way as above. We randomly selected 20 discrete, representative apical myosin puncta and measured their average intensity (Int_p_). Then the number of pixels that have an intensity value larger than Int_p_ was quantified as a proxy for the total amount of apical myosin puncta.

## Statistics

Statistical comparisons were performed using two-tailed Student’s t-tests. Sample sizes can be found in figure legends. *p* values were calculated using MATLAB ttest2 function (Two-tailed Student’s t-test).

## Acknowledgement

We thank Melissa Wang and Amanda Socha for their help with the yeast two-hybrid experiments. We warmly thank Ann M. Lavanway for her help with Spinning Disk Microscope imaging. We thank E. Wieschaus, T. Schupbach, A. Wilde, and C. Field for sharing reagents. We thank the Bloomington *Drosophila* Stock Center (NIH P40OD018537) for providing fly stocks used in this study. We thank all members of the He lab and Griffin lab for helpful discussions. This research is supported by NIGMS ESI-MIRA R35GM128745 and the start-up fund to B.H.

## Author contributions

B. H. and J.C designed the study. J.C. performed the yeast two-hybrid screen. J.C. and B.H. performed the rest of the experiments, analyzed the data, and wrote the manuscript.

## Notes

### Competing Interest Statement

The authors have declared no competing interest.

