## Supplemental Figure 1 - 3 for "Early zygotic gene product Dunk interacts with anillin to regulate Myosin II during *Drosophila* cleavage"

Jiayang Chen and Bing He*

Department of Biological Sciences, Dartmouth College, Hanover, NH, 03755, USA

#### Supplementary Materials

##### Supplementary Figure 1 – 3

### Supplementary Figure 1

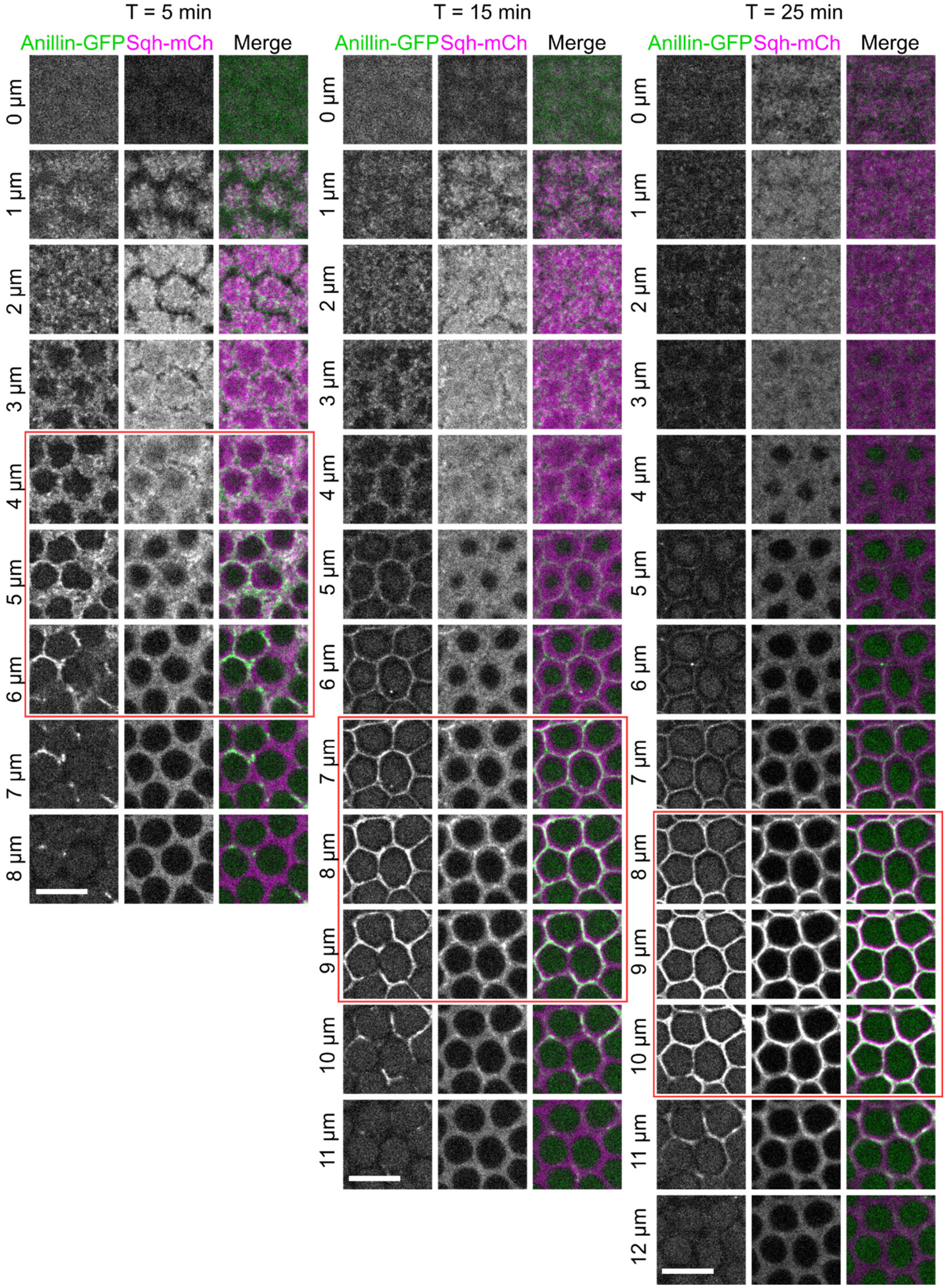


12

**Supplementary Figure 1. Localization of anillin and myosin during early and mid-cellularization.** Live imaging of wild-type embryos expressing anillin-GFP and Sqh-mCherry. Confocal images showing the localization of anillin-GFP and Sqh-mCherry at different z-planes at selected timepoints during early to mid-cellularization. The intensities of the images have been individually adjusted to better demonstrate the co-localization of the two proteins. Red boxes mark the position of the furrow canals. Scale bars: 10 µm**.**

### Supplementary Figure 2

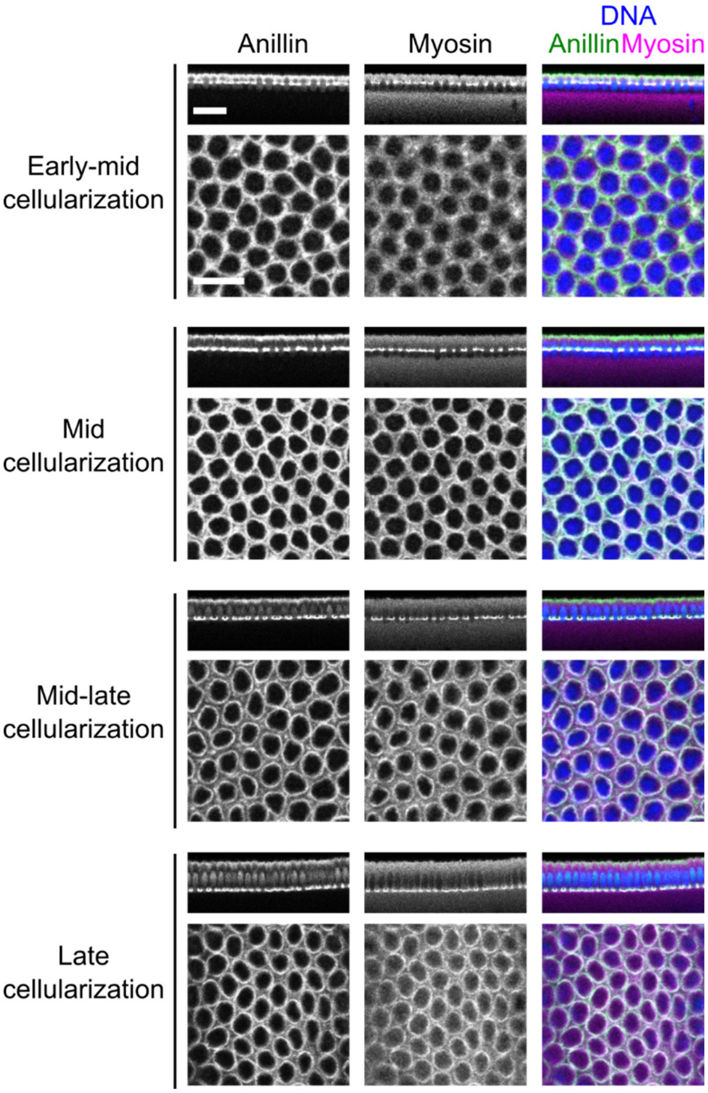


**Supplementary Figure 2. Immunostaining for anillin and zipper (the heavy chain of non-muscle myosin II) in wildtype embryos at different stages during cellularization.** For each stage, top panels: cross-section views showing the stage of the cellularization; bottom panels: en face confocal sections at the level of the furrow canals. Scale bars: Cross-section view: 20 µm; En face view: 10 µm.

### Supplementory Figure 3

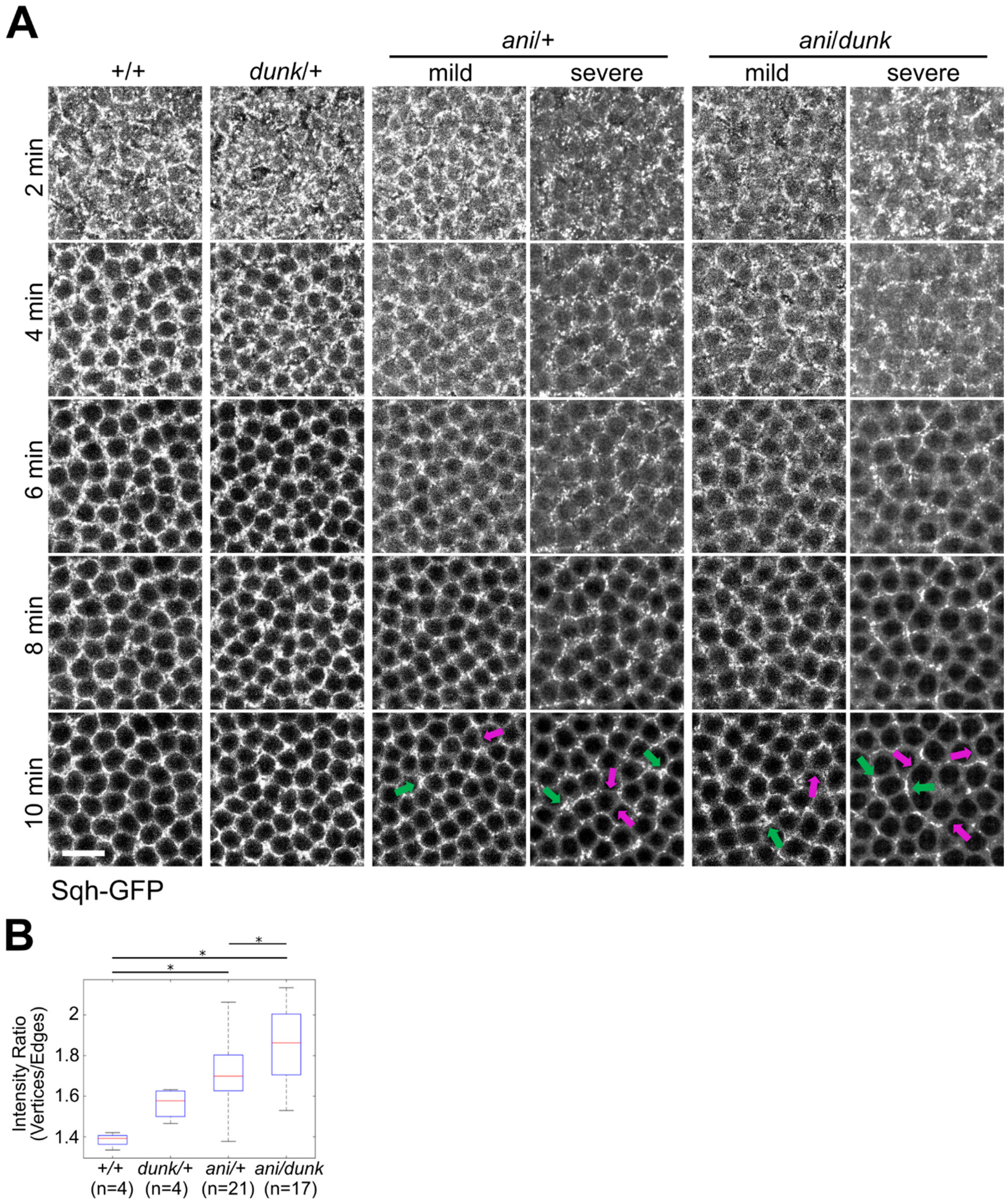

**Supplementary Figure 3. Genetic interaction between *dunk* and *anillin* during early cellularization. (A)** Basal myosin organization in wild type (+/+), *dunk* heterozygous mutant (*dunk/*+), *anillin* heterozygous mutant (*ani/*+) and *anillin dunk* double heterozygous mutant (*ani/dunk*). '*anillin*' here refers to the maternal genotype and '*dunk*' here refers to the zygotic genotype. Shown are maximum projections of ~3 µm confocal sections showing Sqh-GFP at the furrow canals. Basal myosin array in *dunk*/+ embryos was comparable to that in *wildtype* embryos. The distribution of myosin in the basal array is on average more heterogenous in *ani*/+ and *anillin*/*dunk* embryos, with a more biased enrichment at the vertices (green arrow) and depletion from the edges (magenta arrow). The biased distribution of myosin was more prominent in *anillin*/*dunk* embryos than in *anillin*/+ embryos despite the presence of embryo-to-embryo variation in both genotypes (from mild to severe). Scale bar: 10 µm. **(B)** The ratio between vertex- and edge-myosin intensities at 10 min for each genotype. Error bars: s.d. Two-tailed unpaired student t-test was used for statistical comparison. *: $p < 0.05$.
